# Learning gene interactions and functional landscapes from entire bacterial proteomes

**DOI:** 10.1101/2025.03.19.644232

**Authors:** Palash Sethi, Marc G Chevrette, Juannan Zhou

## Abstract

Unraveling complex gene interactions and understanding their functions in the genomes of bacteria will provide critical advancements in fields including bacterial genome evolution, microbiome studies, as well as drug and natural product discovery. This is a challenging problem due to the structural and functional complexity of bacterial genomes, and issues including poor gene annotation in non-model species. Language models (LMs) provide a feasible framework for learning the complex interactions among genes from a large number of publicly available, unannotated bacterial genomes. However, applications of language models have mostly been limited to developing models trained on short genomic sequences. Here, we introduce the first whole bacteria proteome foundation model to our knowledge. Our model was trained on ESM embeddings of tens of thousands of full-size proteomes and can generate contextual embeddings for individual proteins as well as embeddings representing the entire genome. We show that our model captures gene-gene interactions and genomic integrity. We further demonstrate that the learned embeddings can be used to achieve state-of-the-art performances for downstream tasks such as identifying operons, and predicting genotype-phenotype maps.

## Introduction

Genes in an organism interact in complex ways to carry out various metabolic, chemical, physiological, and ecological functions. Frameworks for characterizing gene interactions and the functional landscapes of an organism based on its genomic content have many significant applications in fields including evolutionary biology, genomics, biomedical engineering, synthetic biology, and ecological studies.

For instance, it has been shown that one can achieve better functional annotation for proteins by embedding the proteins of interest in the broad genomic context [1–3], instead of treating them as isolated entities. As another example, several recent studies have characterized natural or synthetic bacterial communities in terms of their ecological productivities. Researchers have also studied the impact of different gut microbiomes on the host’s phenotypes [4–7], such as physiological traits and disease risks. Effectiveness in utilizing this data to predict the function of novel bacterial communities suffer from the limitation that they have largely ignored the functional aspects of the constituent species and mostly treated them as discrete entities. Thus, comprehensive functional characterizations of the species may help us generalize from empirical measurements and make predictions for novel communities by accounting for factors such as convergence evolution, niche partitioning, and cross-feeding.

Generating meaningful representations for genomes that encode gene interactions and diverse functional aspects is a challenging problem. Causes include poor annotations of most non-model organisms and limited success of computational predictions of gene interaction networks. Recent advances in unsupervised models, such as protein language models (LMs) highlight the potential of learning biologically rich information from large amounts of publicly available, unannotated data [8, 9]. However, most published genomic language models (gLMs) are not suitable for providing high-level characterizations of the genome as the majority of them only work on relatively local levels (e.g. short nucleotide sequences) [10–13].

In addition to training gLMs based on raw genomic nucleotide sequences, researchers have also recently started developing LMs on the gene level. For instance, the gLM model in [2] was trained on ESM2 [14] embeddings for individual proteins on the contig level, spanning *∼*30 genes. Although the authors have demonstrated the utility of the model in various applications including improved protein function annotation, it inherently cannot capture long range gene interactions. The local scale also makes it insufficient to provide functional embeddings for the whole genome.

Here, we present BacPT—Bacteria Proteome Transformer—a foundation model specifically designed to perform inferences at both the whole-genome and organism levels. Our model was trained on tens of thousands of full-size bacteria proteomes to reconstruct corrupted protein ESM2 embeddings. It is also able to predict gene families for masked proteins by leveraging the genomic context. Notably, the model captures critical high-level biological information, including diverse forms of gene interactions. Furthermore, we demonstrate that it has learned organism-level functional representations, as evidenced by its superior ability to predict various phenotypes.

## Methods

### Model architecture

We adapt the RoBERTa [15] architecture for masked language modeling tasks in our model BacPT. BacPT has a hidden size of 480, 10 transformer layers with 5 attention heads, and relative key-query position embeddings to capture positional dependencies across sequences of up to 5,000 proteins (Figure 1A). The input for BacPT consist of protein embeddings for all predicted genes predicted by the ESM2 model, esm2 t12 35M UR50D8. Since ESM produces a 480-dimensional vector for each amino acid within a protein, we averaged these vectors across the entire protein to obtain a single representative vector for each gene. A bacterial genome is thus represented by a series of 480-dimensional vectors, each corresponding to a protein. These protein embeddings are then processed through a Multi-Layer Perceptron (MLP) layer and then passed through the transformer layers. Our final model outputs are the predicted protein embeddings of dimension 480.

**Figure 1:**
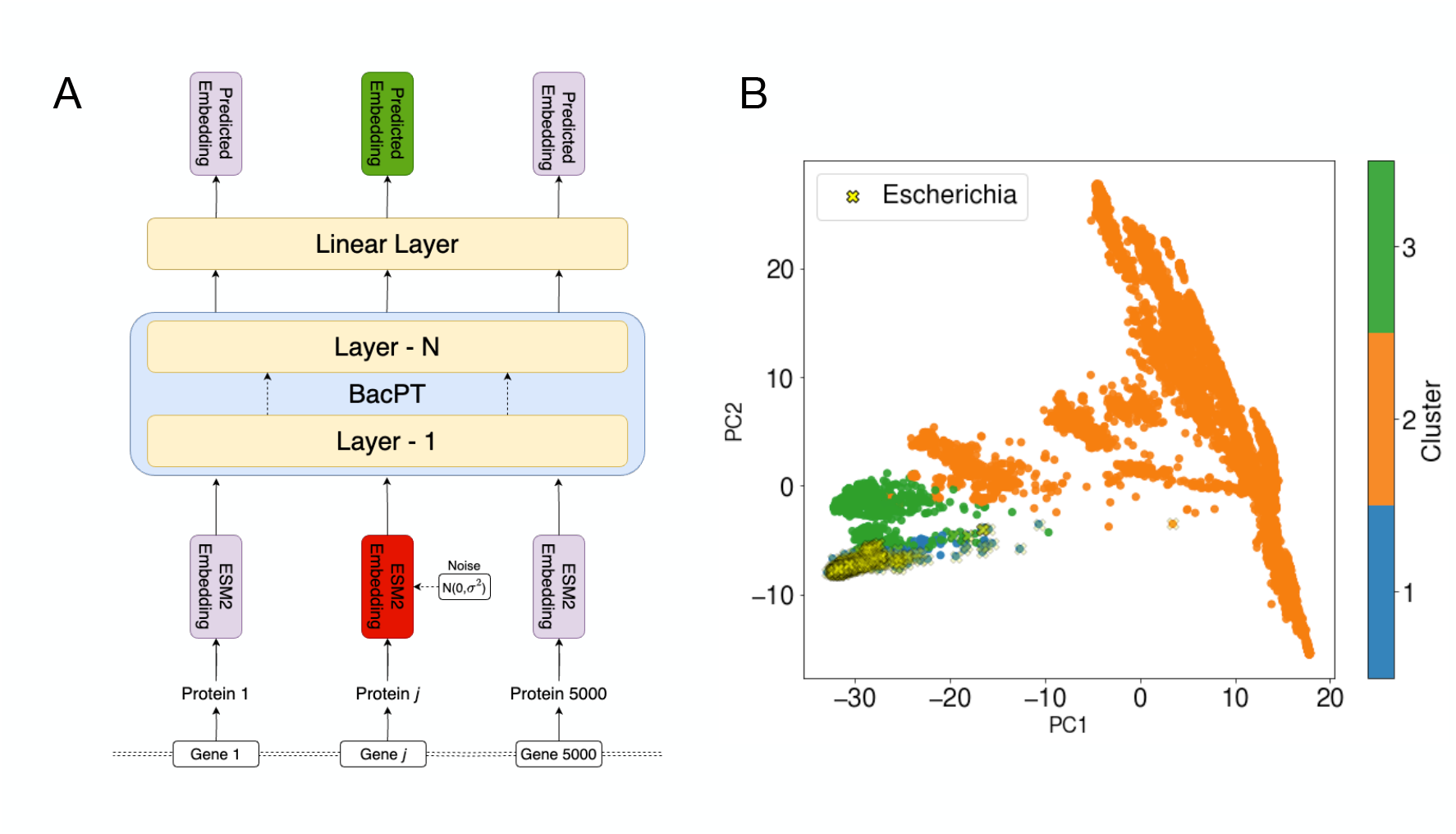
BacPT model architecture and training data. A. BacPT is based on the RoBERTa architecture. Inputs to the model consist of ESM embeddings for all predicted proteins of a given bacterial genome. 40% of proteins are randomly chosen and masked with Gaussian noise. The model was trained on mean square error loss between the original ESM embedding of the predicted embedding. B. Training data was selected based on clustering analysis of protein family representations of 33,140 bacterial genomes. The cluster including the genus Escherichia (blue) was chosen as the test set. Final training data consists of 28,133 genomes.

### Data preparation

Our data consists of protein sequences from the proteomes of 33,140 bacterial species, downloaded from NCBI Genome database [16], representing a diverse array of taxonomic families. To ensure minimal similarity between the genomes in the training and test sets, we conducted a clustering analysis, where we first clustered the proteins across all genomes using MMseqs2 [17] to identify protein families at 50% sequence identity. Each genome was then represented by a vector denoting the presence or absence of each protein family. Finally, we clustered the genomes using HDBSCAN [18] based on their binary representations to elucidate genomic similarities (Figure 1B). Since HDBSCAN is a noise-aware clustering algorithm, we removed genomes classified as noise from our dataset. We selected the distinct cluster containing the genera *Escherichia, Shigella* and *Salmonella* as our test set (consisting of 5059 genomes) and used the remaining clusters for training (92.3 million protein sequences with possible redundancy).

### Model training

We pretrained BacPT using a strategy similar to masked language modeling. Since our inputs comprise continuous embeddings rather than discrete tokens, we progressively corrupted the targeted embeddings with random noise instead of replacing it with a constant vector. Specifically, we applied noise to 40% of proteins randomly selected from a genome and trained the model to reconstruct the original embeddings using mean squared error (MSE) as the loss function. We keep the fraction of noised proteins at 40%, since it has been shown to outperform the masking fraction 15% for large-size BERT models [19]. As training progressed and the noise magnitude was increased in the training set until the masked proteins are completely replaced by Gaussian noise. We hypothesize that BacPT’s ability to reconstruct the raw embeddings of the noised positions from the context of unmasked proteins indicates successful learning of the complex relationships between proteins. We pre-trained the BacPT model for 1100 epochs over 12 days on 16 NVIDIA A100 GPUs.

## Results

### BacPT performance metrics

We validate BacPT’s ability to learn meaningful contextual representations using three complementary approaches. First, we assess whether the model can reconstruct missing protein embeddings solely from surrounding context. Second, we evaluate whether BacPT relies on contextual information rather than memorizing frequently occurring protein embeddings. Third, we analyze whether BacPT embeddings preserve functional relationships by clustering masked proteins and assessing their alignment with predefined gene clusters. These analyses provide insight into how BacPT integrates proteomic context to generate biologically relevant representations.

First, we randomly select 100 genomes from the test set and replace the ESM2 input embeddings of 40% of proteins with Gaussian noise. We then generate BacPT outputs for the noised proteins and compute the Pearson *r*^2^ between these outputs and their original ESM2 embeddings. The mean Pearson *r*^2^ is first calculated for each genome and then averaged across all 100 test genomes. This is benchmarked against unmasked genomes to assess whether the model can accurately reconstruct the missing information. We observe that after a few pre-training epochs, the mean Pearson *r*^2^ for unmasked proteins converges, while the mean Pearson *r*^2^ for masked proteins continues to increase (Figure 2A). This suggests that BacPT can reconstruct protein embeddings solely from the surrounding context, despite the masked protein’s own embedding being entirely absent from the input proteome. Additionally, in the final training epochs, we observe that the masked Pearson *r*^2^ exceeds the unmasked Pearson *r*^2^, indicating that during the forward pass, the model increasingly relies on contextual information rather than the protein’s own embedding.

**Figure 2:**
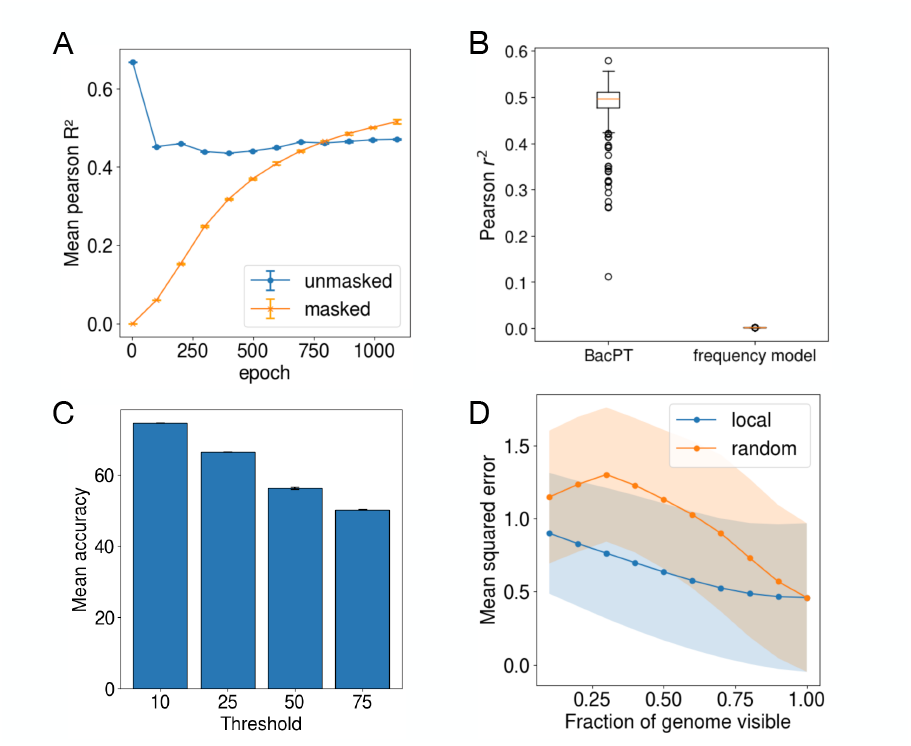
BacPT model performance. A. Correlation between predicted protein embeddings and ground truth ESM embeddings for unmasked and masked proteins on test genomes. Correlation decreases for unmasked proteins as a result of contextualization of the input embeddings. Correlation increases for masked proteins as model’s ability to predict missing genes from the genomic context improves with training. B. Comparison between predictive performance on masked proteins on test genomes for the trained BacPT model and frequency-based model. C. Accuracy of BacPT at predicting the correct gene cluster for masked proteins in test genomes. Results are based on 10,000 gene clusters identified in the training data. Accuracy scores are shown for masked proteins chosen based on different thresholds of Euclidean distance to the cluster centroids. D. Predictions for masked proteins improve when larger proportions of genomes are visible. Furthermore, local genomic context is more important than random context.

Secondly, to ensure that the model is not simply learning to predict the most frequently occurring protein ESM2 embeddings from the training data, we benchmark the masked Pearson *r*^2^ against a frequency-based model (Figure 2B). To construct the frequency model, we randomly selected 500 proteomes from the test set and made predictions for the masked protein by sampling embeddings from proteins in the training set. This approach accounts for the possibility that BacPT might rely on frequently occurring ESM2 embeddings rather than learning meaningful contextual representations. We observe that BacPT’s Pearson *r*^2^ is significantly higher than that of the frequency-based model, indicating that the model captures biologically relevant contextual information rather than memorizing frequent protein embeddings.

Finally, we evaluate our model by clustering the predicted embeddings of masked proteins. While Pearson *r*^2^ provides a useful averaged measure of prediction accuracy, it does not intuitively capture how well BacPT reconstructs masked genes. To address this, we assess whether BacPT can assign masked proteins to predefined gene clusters, leveraging their surrounding proteomic context. The PFAM database [20] groups proteins into tens of thousands of families based on sequence similarity and conserved functional domains, using hidden Markov models (HMMs) to define evolutionary relationships. Following a similar approach, we cluster all proteins in our training data based on their ESM2 embeddings into 10,000 clusters using Faiss-GPU [21]. BacPT embeddings of masked proteins are then mapped to the nearest cluster, and accuracy is defined as the proportion of masked proteins correctly assigned to the same cluster as their ESM2 embeddings. We compute this accuracy for three replicates and evaluate performance across different Euclidean distance thresholds (nearest 10, 25, 50, and 75 percentiles) (Figure 2C). We observe that our model’s prediction accuracy ranges from 74.5 *±* 0.25% at the nearest 10th percentile threshold to 50.2 *±* 0.31% at the 75th percentile, indicating that BacPT embeddings retain strong contextual signals that enable the model predict gene-level information for the masked proteins.

### BacPT captures both local and distant genomic contexts

Next, we investigate whether our genome-scale model has learned to leverage both the full genomic context and the relative importance of local vs. distant genomic information. To assess this, we picked the gene located at the midpoint as the focal gene. We then added Gaussian noise to its embedding and used our trained model to predict the masked embedding while providing protein embeddings for an increasing fraction of genes that surrounds the focal gene or spread randomly across genome. We observed that local genomic context is more informative than random genomic context, as indicated by consistently lower reconstruction MSE when the model had access to embeddings of nearby proteins. Additionally, as more context was provided, the reconstruction accuracy of the focal protein improved (Figure 2D). Notably, even when most of the genome (*>* 90%) was already visible, extending the gene context further enhanced prediction accuracy. This suggests that our model effectively integrates whole-genome context during inference.

### BacPT can identify operons

In this section, we assess if our foundation model has learned correlation between co-occurring genes to enable identification of natural gene clusters. To this end, we focus on identification of operons, which are highly conserved DNA segments containing clusters of adjacent genes that are co-regulated and co-transcribed. We hypothesize that BacPT may capture contextual representations that help distinguish operonic regions from non-operonic regions, as operons exhibit distinct sequence patterns, regulatory signals, and functional associations.

To test this, we curated a dataset consisting of operons containing more than nine genes in the *Escherichia coli* K-12 genome from RegulonDB [22]. To generate non-operonic data for training supervised models, we sampled random genomic contigs from the *Escherichia coli* genome, ensuring their lengths followed a distribution similar to that of the operon data. Additionally, we ensured that these non-operonic regions do not overlap more than 50% with known operonic regions. We generated BacPT embeddings of operon and non-operon data from the last transformer layer of BacPT. We then trained a linear classifier with *L*2 regularization and performed 10-fold cross-validation. To evaluate the effectiveness of BacPT embeddings, we compared them against ESM2 embeddings for operon classification. As shown in Table 1, BacPT achieved an accuracy of 0.75 and an AUROC of 0.83, outperforming ESM2, which achieved an accuracy of 0.66 and an AUROC of 0.75. Our findings first suggest that certain gene families are likely overrepresented in operons, thereby allowing the ESM-based model to make non-trivial predictions. More importantly, they indicate that our model has learned higher-level genomic features beyond individual genes, as reflected in its superior prediction performance. Given the importance of operons in bacterial gene regulation and functional organization, BacPT’s ability to model these structures provides a valuable tool for understanding genomewide transcriptional networks and predicting operon structures in less-characterized bacterial species.

**Table 1:** BacPT identifies operons in *Escherichia coli* K-12 genome. Linear probing of operon embeddings from BacPT outperforms ESM2.

| MODEL | ACCURACY | AUROC |
| --- | --- | --- |
| BACPT | 0.75 | 0.83 |
| ESM2 | 0.66 | 0.75 |

### BacPT learns protein-protein and other gene interactions

Having established that BacPT captures co-transcriptional relationships, we next investigated whether it also learns protein-protein interactions (PPI) and other gene interactions. Specifically, we assessed if our model has learn gene interactions using two approaches: (1) analyzing the attention matrices of BacPT and computing a modified categorical Jacobian by in-silico mutagenesis. We performed both experiments on *Escherichia coli* K-12 genome.

First, to evaluate whether BacPT’s attention weights correspond to PPI strength, we performed a correlation analysis between attention weights from all the attention heads and transformer layers of BacPT against an experimental PPI strength dataset for the *Escherichia coli* K-12 genome [23]. The attention weights were obtained by passing the *Escherichia coli* K-12 genome through BacPT without masking. To refine the interaction signal, we symmetrized the attention matrices, subtracted the average product correlation, and retained the attention weights corresponding gene pairs present the PPI dataset. We inferred the relationship between the attention weights of BacPT and the experimental PPIs by calculating the Pearson correlation, which is compared against a null distribution of Pearson correlation values generated using attention weights of randomly initialized models. Overall, we observed weak to moderate correlations across all layers and attention heads, with *p* values ranging from nonsignificant to highly significant. The most significant correlation was observed for 2nd attention head in the 7th layer (*r* = 0.15, *p* = 10^*−*18^) (Figure 3B). Additionally, the correlation between the attention weights of the trained model and the ground truth is systematically higher than the null distribution. This shows that BacPT’s attention mechanisms capture biologically relevant interaction patterns, with certain layers and heads specifically encoding protein-protein interaction signals.

**Figure 3:**
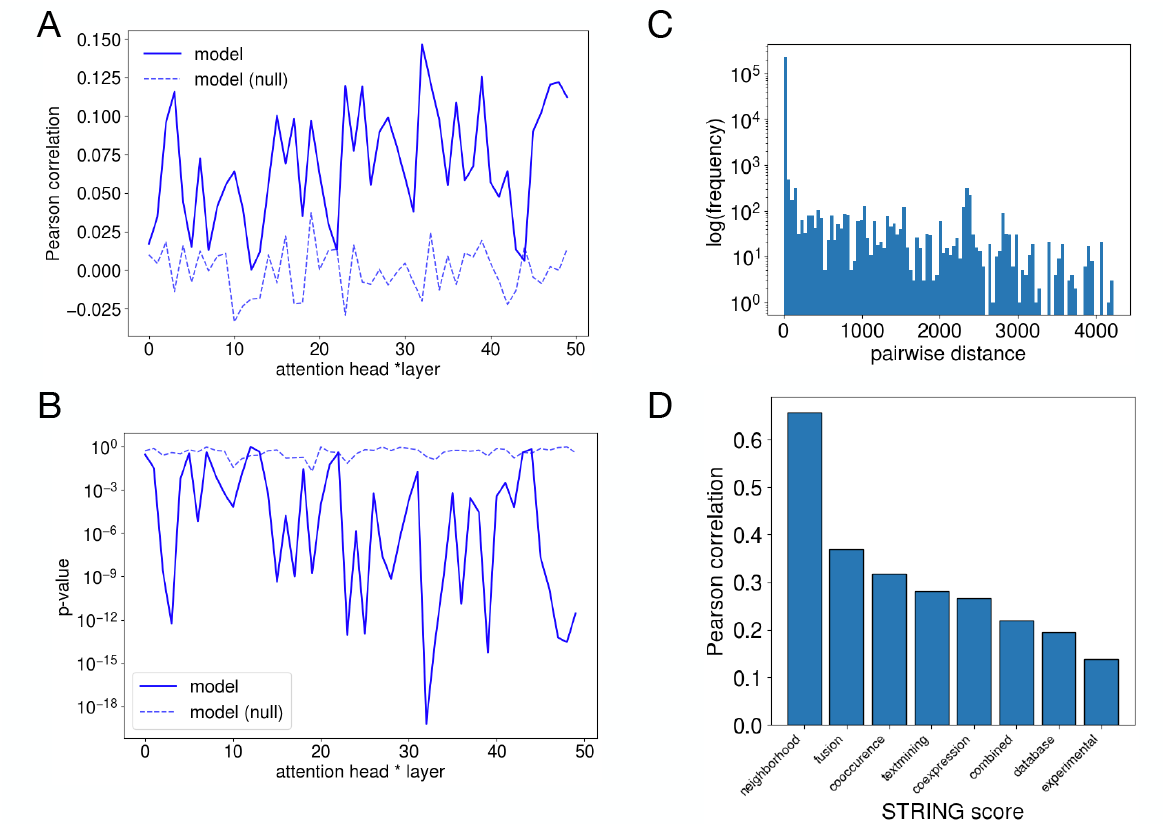
BacPT captures protein-protein interactions (PPI) as well as other gene interactions. A. Pearson correlation between attention weights for each attention head and transformer layer of trained BacPT model and the experimental PPI dataset from [23]. B. P-values for the correlation. C. BacPT captures both local and long-range interactions. Histogram shows distribution of distances in terms of genes for the top 99% pairwise interactions. D. Pearson correlation between pairwise interaction coefficients estimated as the Jacobian of the trained model and various types of gene-gene interactions extracted from the STRING database.

We next extracted a single matrix from our trained model to summarize the overall gene relationship predicted by our model and compared this matrix to the ground-truth PPIs. Our method is similar to the categorical Jacobian [24], which quantifies pairwise interaction between amino acids by introducing categorical perturbations to the ESM2 input sequence by replacing one amino acid token at a time. However, since BacPT operates on protein embeddings rather than discrete tokens, it lacks a categorical perturbation vocabulary. To adapt this method, we replaced each input protein embedding with a zero vector and measured the resulting changes in BacPT’s output across the entire proteome. This provides an estimate of the extent to which the perturbed protein influences the predictions for other proteins. We then computed the influence of the focal masked gene *i* on gene *j* as the mean squared difference between the BacPT predictions for *j* made with and without masking *i*. This produced an *L × L* interaction matrix for a genome of length *L*. Finally, we symmetrized the matrix and subtracted the average product correlation to refine the interaction signal.

We first observed that the distribution of Jacobian scores is highly skewed, with most gene pairs having low scores and a small subset exhibiting significantly higher values, potentially indicating strong gene interactions (Figure S1-Appendix). We next examined the distribution of distance between genes among the top 1% strongest interactions, where distance was calculated as differences in position indices. We see that although most interacting genes are physically close, our model revealed strong interactions over long physical distances, spanning up to thousands of genes (Figure 3C). This result suggests that BacPT does not solely rely on genomic proximity but also encodes distant associations, suggesting an ability to model regulatory and structural dependencies beyond immediate neighbors. We tested this hypothesis by comparing Jacobian weights with various scores from the STRING database [25]. Specifically, we compared the entries in our Jacobian matrix against various STRING gene interaction values. While STRING scores do not directly measure interaction strength, they represent probabilities derived from multiple evidence channels while accounting for the likelihood of randomly observing an interaction. The STRING scores we compared against include channels corresponding to neighborhood, fusion, co-occurrence, text mining, coexpression, database, experimental, as well as a combined score. Thus, significant correlation between the Jacobian matrix and a type of STRING scores indicates that our model has learned certain aspects of gene interactions. We included only interactions where the combined score meets a minimum threshold of 0.9 to ensure a high-confidence set of interactions, in order to the reduce the influence of spurious or low-confidence associations. We found that our Jacobian (Figure 3D) has the highest correlation with neighborhood-based interactions (Pearson *r* = 0.65), followed by fusion and co-occurrence (Pearson *r* = 0.37 and 0.31). This suggests that BacPT effectively captures genomic proximity and evolutionary signals. This is expected as we have found earlier that although our model is able to model the whole proteome, it preferentially leverages local-level information. Notably, we observed a level of correlation (*r* = 0.15) between the Jacobian and STRING scores corresponding to experimentally measured protein interactions, similar to the strongest correlation observed between attention weights and the K-12 PPI datasets.

### BacPT representations enable enhanced prediction of organism-level traits

Functional characterization of bacteria at the organism level based on genomic data has significant applications including phenotype prediction and the study of microbial community interactions. We hypothesize that BacPT, by contextualizing genes through interactions across the whole proteome, provides functionally rich embeddings that can be exploited to improve the prediction accuracy across different categories of bacterial traits.

To test this, we curated a genome-phenotype dataset using Bac*Dive* [26], a database of standardized prokaryotic strain-level data. We extracted 99 traits for 5059 test proteomes from Bac*Dive*, spanning a broad range of functional and phenotypic characteristics, including enzyme activity, metabolite production, metabolite utilization, and morphological properties. For each proteome in BacPT’s test set, we extracted a 480-dimensional vector from each attention layer by averaging over the embeddings across all proteins. For each trait extracted from Bac*Dive*, we train 10 linear probes, each based on embeddings from one of the 10 attention layers, to map BacPT’s embeddings to Bac*Dive* traits under *L*2 regularization, with regularization parameter chosen by five-fold cross-validation. We performed 10 separate logistic regressions in order to preserve the most amount of biological information since we do not know a priori which layers produce the most relevant embeddings for a given phenotype. We evaluated performance of the linear probes using *F*1 score and report the best *F*1 score across 10 models for each trait.

To benchmark the performance of our method, we also fit Traitar [27], a framework for predicting a list of predefined traits from only the genome sequence. Traitar operates by identifying relevant protein families from the training data and leveraging their presence to predict traits. Additionally, we also fit linear probes using the mean protein ESM embeddings.

We first compared the performance of the three models using the *F*1 score for 30 out of 99 total Bac*Dive* traits, for which Traitar is able to make predictions (Figure 4A). Out of the 30 traits, BacPT outperforms Traitar in 22 traits and achieves comparable results in 5 traits. We next compared the performance between the BacPT-based linear probes and the ESM2-based linear probes on all 99 traits. Given that the ESM2-based linear probes can only model the independent contribution of genes, this comparison is relevant for assessing how incorporation of contextualized gene embeddings and gene interactions can help the functional characterization on the whole organism level. Overall, we found that BacPT embedding almost always increase prediction performance (Figure 4B). Importantly, we observed an average 5% improvement in F1 score with BacPT, and substantial improvement (Δ F1 score *>* 0.2) for certain traits (e.g. Lipase activity). Our results demonstrate that BacPT effectively captures biologically meaningful features in bacterial proteomes, which can enable superior trait prediction compared to both Traitar and ESM2 embeddings. The improved performance highlights the ability of BacPT’s contextualized embeddings to encode functional and phenotypic characteristics, suggesting its potential as a robust tool for large-scale microbial trait prediction. This approach provides a scalable and data-driven method for linking genotypic information to phenotypic traits, which could enhance our understanding of bacterial adaptation, metabolic capabilities, and ecological interactions.

**Figure 4:**
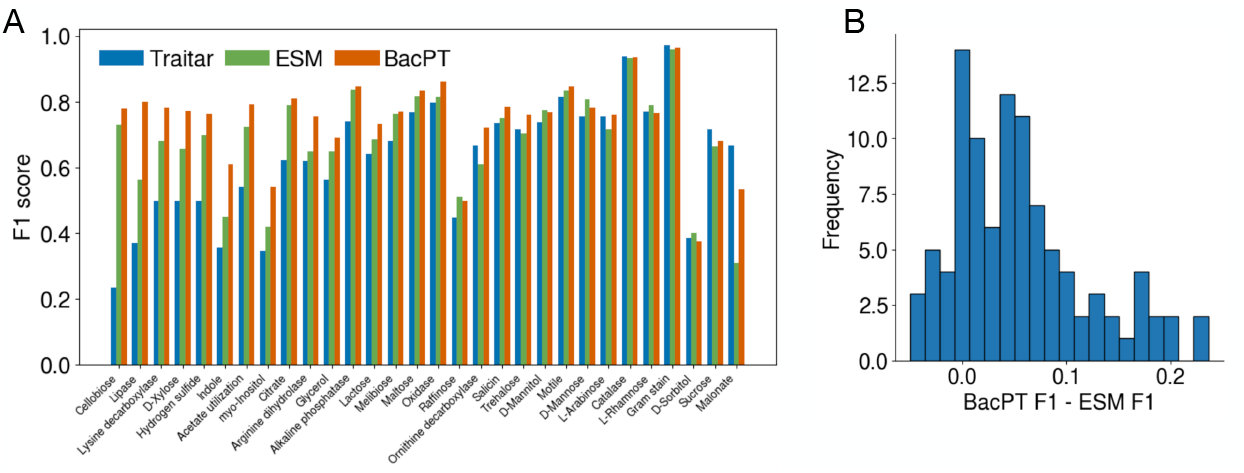
BacPT representations enable enhanced prediction of organism-level traits. A. *F*1 score for Traitar, a standard method for predicting phenotypes based on whole genome sequences, as well as linear probes trained on mean ESM embeddings and BacPT embeddings. BacPT embeddings from the final and intermediate layers were used to train individual linear probes. Performance of the best probe is shown for each trait. B. Performance improvement of BacPT compared with ESM. The difference in *F*1 value between the best linear probe for BacPT and the ESM linear probe was calculated for 99 traits derived from the Bac*Dive* database.

## Discussion

In recent years, the significant reduction in sequencing costs has led to the accumulation of vast amounts of whole genome sequence data. Substantial efforts have been made to extract biological insights from these natural genomes. However, to date, nearly all genomic foundation models have been limited to much smaller scales compared to the entirety of whole genomes. Here, we present, to our knowledge, the one of the first foundation model trained on whole genomes. Using mean ESM embeddings for individual proteins in place of tokens, we successfully trained a transformer model capable of modeling interactions among all genes (*∼* 5, 000) within a bacterial genome.

We demonstrated that our model can accurately predict masked genes based on their genomic context. Additionally, we showed that the model effectively leverages both local and global genomic contexts for its predictions. Importantly, we found that our model captures diverse types of gene interactions and the latent embeddings of the whole genome provide rich biological representations, useful for making accurate predictions of phenotypic traits at the whole-organism level.

Our trained model exhibits the strongest correlation with the ‘neighborhood’ associations between proteins [25]. This is not surprising, since our experiment showed that our model preferentially uses local gene context for making predictions. This suggests that our model has effectively captured signals of co-occurrence among proteins that are frequently organized into gene clusters across diverse taxa. Thus, our model holds potential utility for identifying prevalent yet unannotated gene clusters, including novel biosynthetic gene clusters.

Interestingly, our model also exhibits moderate yet significant correlations with other types of gene associations, such as protein-protein interactions and co-expression. This supports the hypothesis that gene interactions can be inferred from patterns of gene covariation, analogous to how protein structure can be predicted from couplings between conserved amino acids. However, unlike proteins, gene co-occurrence is influenced by multitude of biological factors. As a result, future supervised frameworks could be developed to specifically identify protein-protein interactions based on our pretrained model.

In addition to discovering gene interactions, our model also provide an unsupervised framework for functionally characterizing whole genomes. We demonstrate this utility by using the latent embeddings of our model for predicting organism-level metabolic, fitness, and morphological traits. We observed that our model frequently outperforms the standard Traitar method, as well as mean protein embeddings. This may be attributed to the model’s ability to contextualize causal genes or capture key gene interactions. Further applications of BacPT’s embeddings could include predicting novel traits in uncharacterized bacteria, refining genotype-phenotype association studies, and aiding in the functional annotation of microbiomes. Overall, this result suggests that foundation models trained at the whole-genome level hold significant promise for many downstream applications requiring rich organism-level representations.

## Appendix

### Figures

**Figure S1:**
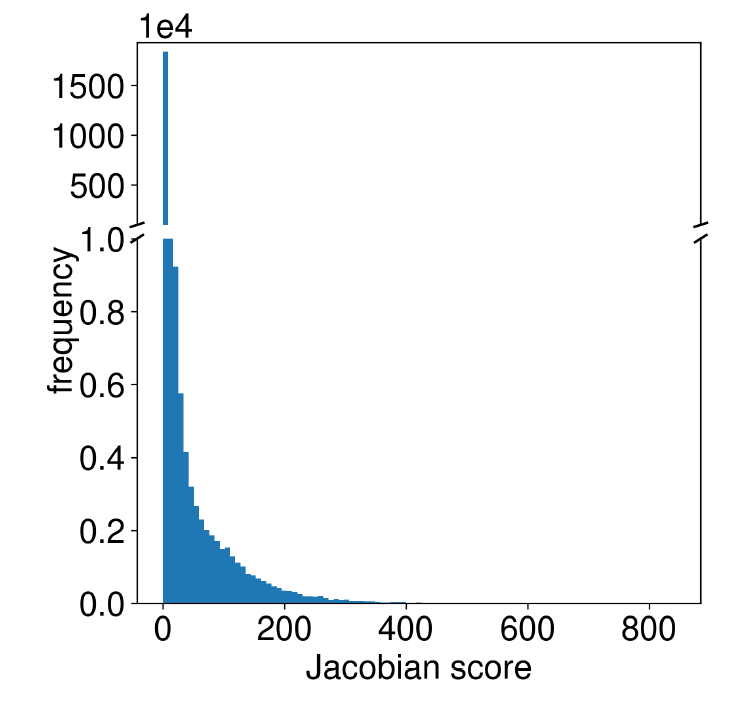
Distribution of Jacobian scores for pairs of genes in the test E. coli K-12 genome.

## Notes

### Competing Interest Statement

The authors have declared no competing interest.

